# An ImageJ-based tool for three-dimensional registration between different types of microscopic images

**DOI:** 10.1101/2022.09.11.507445

**Authors:** Hiroshi Koyama, Kanae Kishi, Toshihiko Fujimori

## Abstract

Three-dimensional (3D) registration (i.e. alignment) between two microscopic images is substantially helpful to study tissues not adhere to substrates such as mouse embryos and organoids which are often three-dimensionally rotated during imaging. However, there is no 3D registration tool easily accessible for experimental biologists. Here we developed an ImageJ-based tool which achieves 3D registration accompanying both quantitative evaluation of the accuracy and reconstruction of 3D rotated images. In this tool, several landmarks are manually provided in two images to be aligned, and 3D rotation is computed so that the distances between the paired landmarks from the two images are minimized. By simultaneously providing multiple points (e.g. all nuclei in the regions of interest) other than the landmarks in the two images, the correspondence of each point between the two images is quantitatively explored: a certain nucleus in one image corresponds to which nucleus in another image. Furthermore, the 3D rotation is applied to one of the two images, resulting in reconstruction of 3D rotated images. We demonstrated that this tool successfully achieved 3D registration and reconstruction of images in mouse pre-implantation embryos, where one image was obtained during live imaging and another image from fixed embryos after live imaging. This approach provides a versatile tool applicable for various tissues and species.

## Introduction

Three-dimensional (3D) imaging is a central technique in developmental biology and organoid studies, which is achieved by confocal microscopies, multiphoton microscopies, micro-CT (computed tomography), etc. During live imaging of tissues such as early embryos of mice, chordates, and echinoderms, and organoids, they can be three-dimensionally rotated in the cultured medium/liquid because they do not adhere to substrates. In studies relating to cell tracking and cell lineage, researchers have to pay much efforts to determine the correspondence of each cell between images before and after the rotation (Koyama et al., 2022; Kurotaki et al., 2007). Similar issues also raise when researchers compare live imaging data with images obtained from samples fixed after live imaging (Fig. 1) (Pokrass et al., 2020; Simon et al., 2020); e.g. characteristics different from those obtained from the live imaging are visualized from fixed samples that are subjected to immunostaining, while the tissues are rotated during fixation. In addition, alignment between different embryos at a similar embryonic stage is also helpful for comparing differences of cell lineage and positions between the different embryos (Onuma et al., 2020). Under these situations, researchers physically correct the rotations during sample preparations on microscopic stages or correct on the basis of manual image processing (Onuma et al., 2020; Pokrass and Regot, 2021; Simon et al., 2020), both of which are usually time-consuming processes. In the former case, an exact correction is almost impossible (i.e. spatial discrepancies between two images remain to some extent), which may be problematic for spatially intricate regions in tissues. In the latter case, the correspondence of each cell between two images is performed without reconstructing three-dimensionally rotated images, and thus the outcomes of the operations are not clearly presented. Therefore, it is difficult for other researchers (and even for the researchers doing the manual operations) to evaluate whether the corrections are reliable. To improve these situations, it is critical to develop an image processing tool for 3D registration accompanying both visualization and quantitative evaluation of the outcomes, which should be easily accessible for experimental biologists who are not so familiar to image processing.

**Figure 1:**
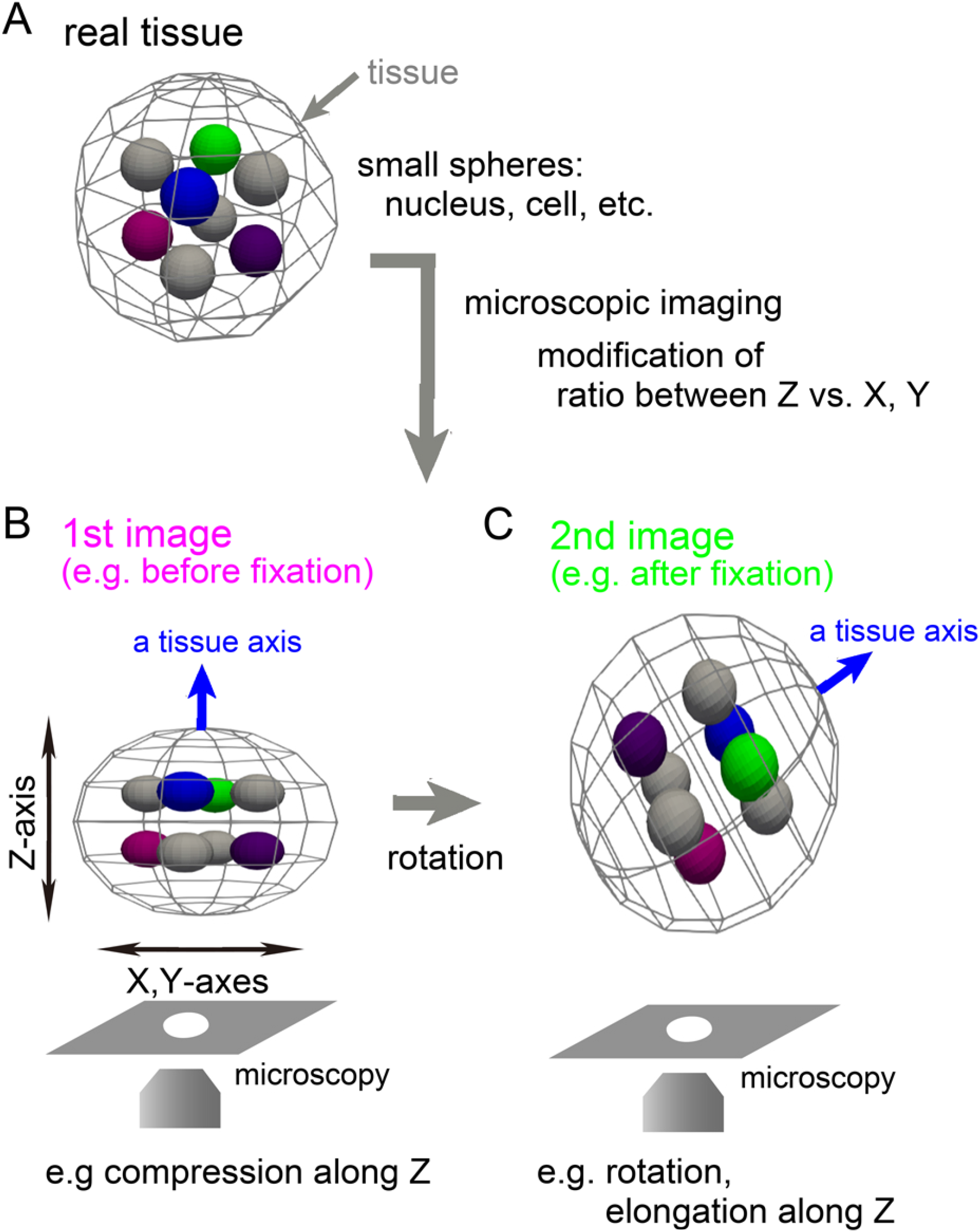
Illustration of rotation and distortion of specimen during preparation A. A tissue is illustrated with inner objects. In this case, the tissue and the inner objects are depicted as spheres. B. An example of image acquisition of the tissue is shown. The acquired image can be shrunk or elongated along Z-axis. C. A rotated tissue is shown. During experimental procedures including fixation, the tissue may be rotated (“a tissue axis” between B and C).

For experimental biologists, the ImageJ software and its high-functionality version Fiji are the most widely accessible image processing tools. In default plugins implemented in Fiji, “Correct 3D drift” is a 3D registration tool, but it only considers 3D translation (i.e. x, y, and z-directional movements) but not rotation. Another 3D registration plugin called “Descriptor-based registration (2d/3d)” is implemented for specific situations where many beads are embedded as landmarks in samples and are computationally detected for subsequent usage of 3D registration (Preibisch et al., 2010).

A more primitive way is manual operations of “Rotate” and “Reslice” tools, both of which are basic functions in ImageJ/Fiji; through multiple cycles of these two tools, any 3D rotation and subsequent 3D reconstruction can in principle be achieved. However, as far as we tried, it is very hard to determine correct 3D rotations and angles of reslices, probably except for researchers who can easily imagine 3D rotation of objects. Image processing tools other than ImageJ have been developed for experimental biology especially for segmentation of two-dimensional epithelial cells and subsequent quantitative analyses (Heller et al., 2016; Tan et al., 2021), and for segmentation of objects in three-dimensional tissues (Azuma and Onami, 2017; Bao et al., 2006; McDole et al., 2018).

In the present study, we developed a versatile 3D registration/rotation tool which can be applicable for any types of 3D images and can be run on ImageJ.

## Materials and Methods

### Mouse embryos

The mouse embryos at the blastocyst stage were used as the test case. We performed confocal fluorescent microscopic imaging of fluorescently-labeled nuclei in living embryos in a manner similar to our previous works (Fig. 2, 1^st^ image) (Azuma and Onami, 2017; Koyama et al., 2022). Subsequently, we fixed the embryos, stained the nuclei by Hoechst, and then imaged them (Fig. 2, 2^nd^ image). The blastocysts are composed of two cell types: the trophectoderm (TE) cells form an outermost layer and the inner cell mass (ICM) cells form an inner cell aggregate with high cell density (Fig. 2, illustration). The image obtained from the fixed embryos showed rotation compared with the image from living embryos (Fig. 2; see the location of ICMs).

**Figure 2:**
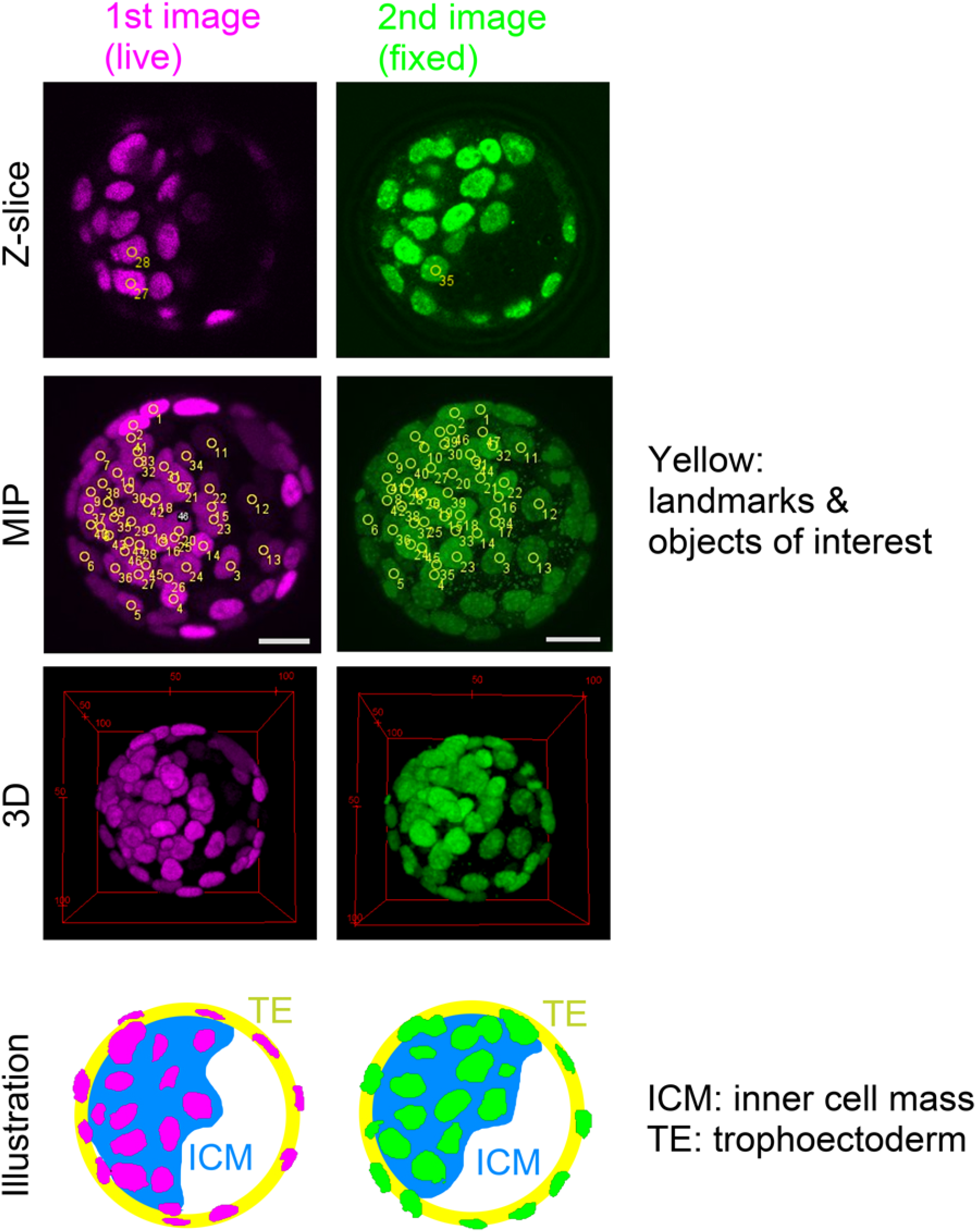
Rotated image of mouse blastocyst The 1^st^ and 2^nd^ images are acquired from a live or fixed embryo. A Z-slice, maximum intensity projection (MIP), and 3D view of the images are shown. The regions of the inner cell mass are illustrated for each image. Landmarks and objects of interest were labeled by using ImageJ>Multi-point tool.

The microscopic imaging conditions are as follows: the confocal microscopy (A1 laser scanning confocal microscope, Nikon, Japan) with a 60× objective (PlanApo; WI; NA=1.20, Nikon, Japan), Z-slices separated by 0.575μm for live embryos or 0.625μm for fixed embryos. The nuclei in the live or fixed embryos were labeled by YFP conjugated with a nuclear localization signal or Hoechst, respectively.

### Overview of methods for 3D-registration

For studies of cell lineage in the mouse early embryos (Kurotaki et al., 2007; Pokrass et al., 2020; Simon et al., 2020), 3D registration during live imaging or between live and fixed embryos substantially supports the analyses. We developed a landmark-based 3D registration and applied to the blastocysts. The overview of our method is illustrated in Fig. 3. In the first step, as landmarks, we manually chose several pairs of the same nuclei between the 1^st^ and 2^nd^ images (Fig. 3A, step-1). In addition to the landmarks, we manually chose all objects (i.e. nuclei) of interest in the both images, which are not paired at this moment (Fig.3A, step-2). The manual steps needed in our method are limited to the above two steps, and thus, user’s effort is minimum. The next step is the core in our method, where 3D rotation was computationally performed so that the summation of the distances between the paired landmarks became minimized (Fig. 3A, step-4). Note that this summation is called the cost function to be minimized. In the present case, the landmarks in the 2^nd^ image were rotated. In general, 3D rotation is expressed as a matrix composed of three rotational angles (Fig. 3B), while 2D rotation is of one rotational angle (Fig. 3B). Therefore, the optimal values of these three angles were computed; the mathematical algorithm is explained in Appendix.

**Figure 3:**
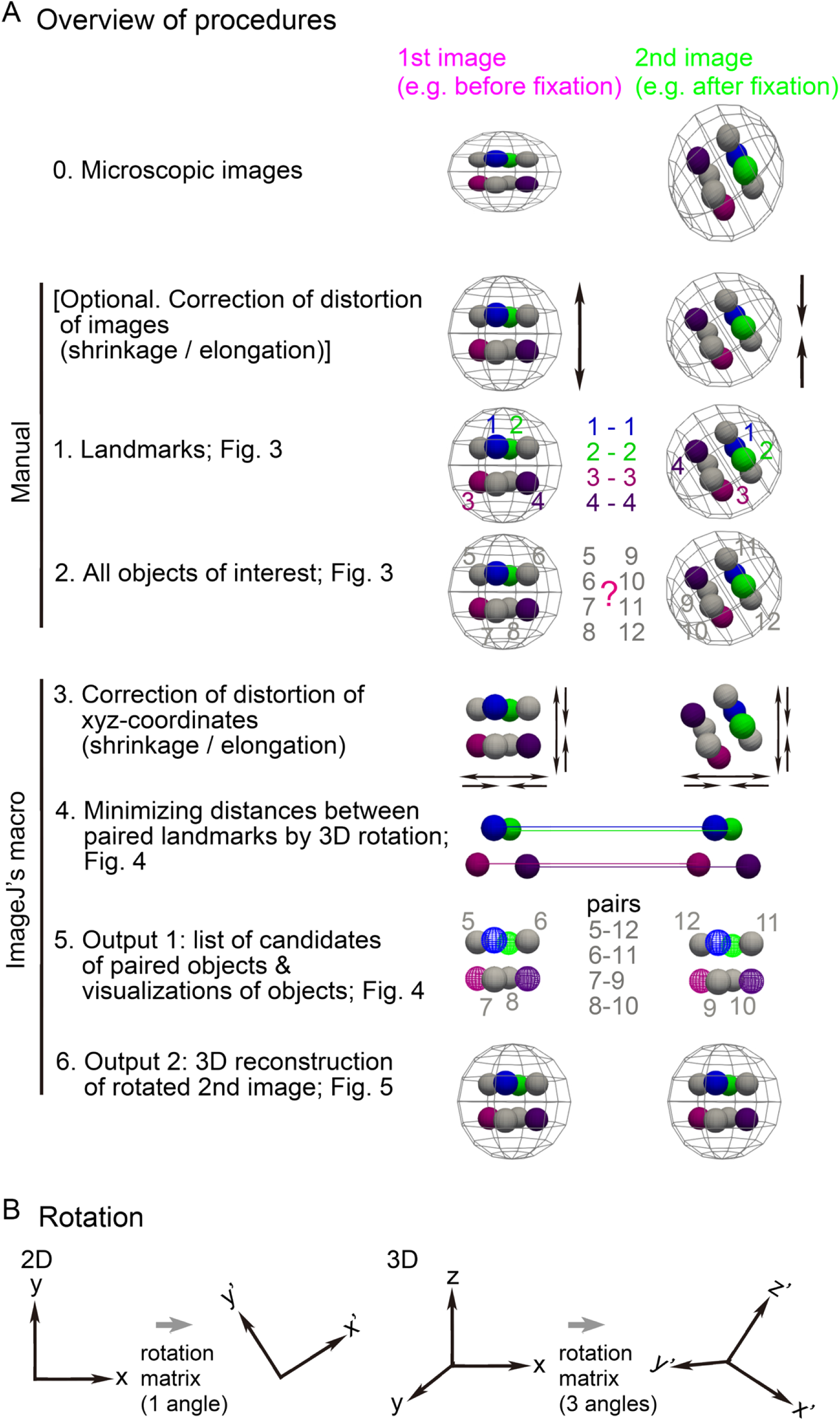
Procedures of 3D registration and reconstruction A. The procedures of our method are illustrated. At step-0, microscopic images are shown where the 2^nd^ image is rotated compared with the 1^st^ image. At the “Optional” step, the shrinkage or elongation of the two images is corrected (arrows). At the step-1, 4 landmarks are exemplified (#1-4). At the step-2, objects of interest are labeled by non-overlapped numbers between the two images (#5-8 vs #9-12). At the step-3, shrinkage or elongation of the xyz-coordinates of the landmarks and the objects of interest are corrected. If shrinkage or elongation of the images has been already corrected at the “Optional” step, the step-3 is not required. At the step-4, the landmarks in the 2^nd^ image are optimally rotated. At the step-5, the paired objects are identified (e.g. 5-12, 6-11). At the step-6, the 2^nd^ image is rotated to be aligned with the 1^st^ image, and the rotated image is reconstructed. B. Definition of 3D rotation is explained. In the case of 2D rotation, the rotation matrix contains one angle (*θ* in Appendix 1). In the case of 3D rotation, the rotation matrix contains three angles (*ϕ, θ*, and *Ψ* in Appendix 1).

By using the optimal values of the three angles, we performed following two analyses. 1) The xyz-coordinates of the nuclei of interest chosen in the previous step were computationally rotated according to the three rotational angles (Fig. 3A, step-5). For each nucleus in the live embryos, we computationally determined the nearest nucleus in the fixed embryos. In other words, we determined the correspondences of the several tens of the nuclei between the live and fixed embryos (Fig. 3A, step-5, “pairs”). 2) We also applied the rotation based on the three rotational angles to the images from the fixed embryos, and reconstructed 3D images (Fig. 3A, step-6). Consequently, we can easily compare the resultant 3D images with the images from the live embryos. These two analyses enabled us to quantitatively and visually determine the correspondences of the nuclei between the live and fixed embryos.

In addition to the above steps, we implemented optional steps to adjust real situations. In real tissues, fixation often shrink tissues. Moreover, conditions of microscopic imaging cause shrinkage or elongation of 3D images along the Z-axis; e.g. differences of refractive indices between glass of the glass-base dishes and medium for specimen. Severe shrinkage or elongation can spoil the 3D registration. We can revise the x, y, and z scales in both the live and fixed embryos before or after the choice of the landmarks (Fig. 3A, “[Optional…” and step-3).

## Results

### 3D rotation of landmarks and objects of interest

We applied our method to the mouse blastocysts. In Fig. 2 (“Z-slice” and “MIP”), we chose 9 nuclei as landmarks. In addition to the landmarks, we also labeled several tens of the nuclei as objects of interest. We loaded the xyz-coordinates of the landmarks into our ImageJ-macro, and then, computationally obtained the values of the three angles for 3D rotation. Note that the rotation centers in the two images were set at the centroids of the landmarks. Finally, we generated an image where the rotated positions of the landmarks were depicted as particles. This image is a 3D image (i.e. composed of multiple z-slices) Fig 4A shows a merged 3D image of the landmarks from the 1^st^ and 2^nd^ images. Before the rotation, the landmarks from the two images were not closely located (Fig. 4A, left panel), whereas, after the rotations, the landmarks became closely located (Fig. 4A, right panel).

**Figure 4:**
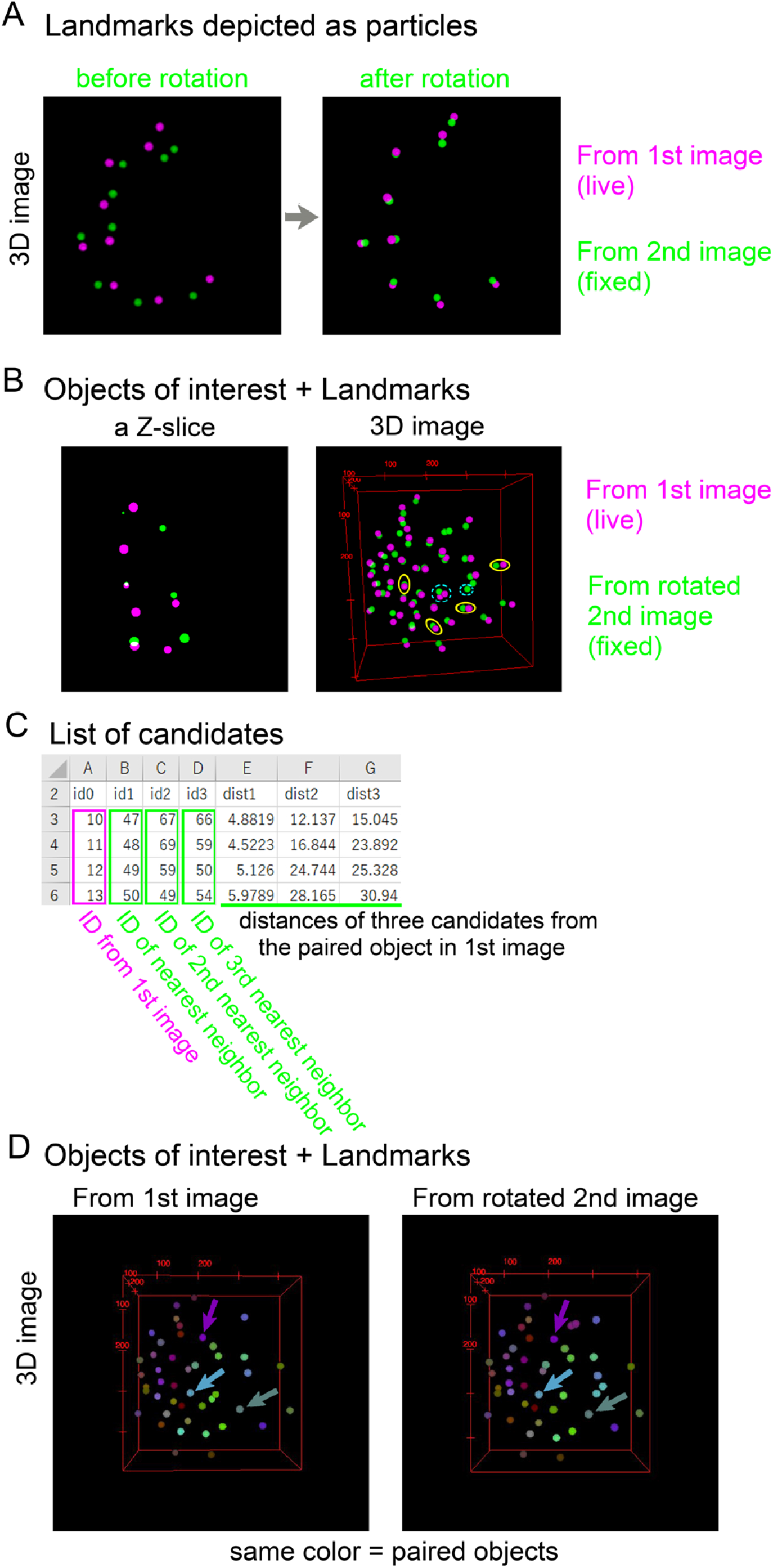
Registration of landmarks and objects of interest A. Landmarks in the 1^st^ and 2^nd^ images are depicted as particles in 3D images. Images before and after the rotation of the 2^nd^ image are shown. The 3D images were generated by using Fiji>Plugins>3D Viewer; all 3D images in this article were generated by the 3D Viewer. B. Objects of interest in the 1^st^ and 2^nd^ images are depicted as particles in 3D images. The landmarks are also depicted. Yellow circles, some examples of paired objects; light blue circles with dashed lines, a few examples of unsuccessfully paired objects. C. Quantitative evaluation of pairing. For each object of interest in the 1^st^ image, three objects as a candidate for pairing are shown in the 2^nd^ image according to distances between the objects. Four objects in the 1^st^ image are exemplified. In the case that an object in the 2^nd^ image is multiply assigned as the nearest neighbor for different objects in the 1^st^ image, such multiply-assigned objects are also listed in the output text file (not shown in this figure). D. Paired objects between the 1^st^ and 2^nd^ image are depicted as particles in the same color. Arrows, three examples of paired objects. Landmarks are also depicted. The original images were 8-bit images where the intensities of each particle correspond to the IDs of the objects, and the color was provided by setting lookup tables (ImageJ>Image>Lookup Tables>3-3-2-RGB).

Then, we applied the 3D rotation to the positions of the nuclei other than the landmarks, and generated a 3D image (i.e. composed of multiple z-slices). In the 3D image where the nuclei from the 1^st^ and the rotated 2^nd^ images were merged (Fig. 4B, left panel), most of the nuclei from the two images were closely located and paired each other (e.g. labeled by yellow). We can also find several nuclei which did not have counterparts (e.g. labeled by light blue with dashed line), this is because we failed to label all the nuclei at the step-2 in Fig. 3. In other words, the 3D visualization helped us to judge whether we successfully label all nuclei of interest. In addition, we can also check the localization of the positions of the rotated nuclei in each Z-slice (Fig. 4B, right panel).

To determine the correspondence of each nucleus between the 1^st^ and the rotated 2^nd^ images, we calculated the distances between the nuclei in the two images. For each nucleus in the 1^st^ image, we searched for the nearest nucleus in the rotated 2^nd^ image. Fig. 4C show the ID of the nearest nucleus and the distance between the paired nuclei. We also searched for the 2^nd^ and 3^rd^ nearest nuclei as shown in Fig. 4C. By quantitatively evaluating the distances of these three candidates, we can judge which nucleus is the counterpart of each nucleus from the 1^st^ image. Simultaneously, under an assumption that the nearest nucleus is the correct counterparts, we generated a 3D image where the paired nuclei were presented by the same color (Fig. 4D).

### 3D reconstruction of rotated image

We developed an algorithm to reconstruct rotated 2^nd^ images. We applied the above 3D rotation to the 2^nd^ image itself (i.e. pixel/voxel-based rotation is applied). In Fig. 5A, the rotated 2^nd^ image is shown (blastocyst #1 vs. Fig. 2 which are the images before the rotation). A merged image of the 1^st^ and the rotated 2^nd^ images exhibited good correspondences of the nuclei. Moreover, the correspondences are also confirmed by visualizing Z-slice images in Fig. 5B (e.g. labeled by yellow). The slight spatial discrepancies between some pairs of the nuclei may result from shrinkage of the blastocyst by the fixation. Chromosome segregation was observed in the 2^nd^ image (Fig. 5B, at the upper right of Z-slice #2, which was stained by Hoechst), whereas the signal in the 1^st^ image was obscure; this is because the nuclear localization signal which does not bind to the chromosomes was used in the 1^st^ image. Another example of the blastocyst is shown in Fig. 5A (blastocyst #2). Although the directions between the 1^st^ and the 2^nd^ image before the rotation were quite different (1^st^ image vs. before rotation), the directions became absolutely aligned after the rotation (1^st^ image vs. rotated 2^nd^ image), and the merged image showed clear correspondences between the paired nuclei.

**Figure 5:**
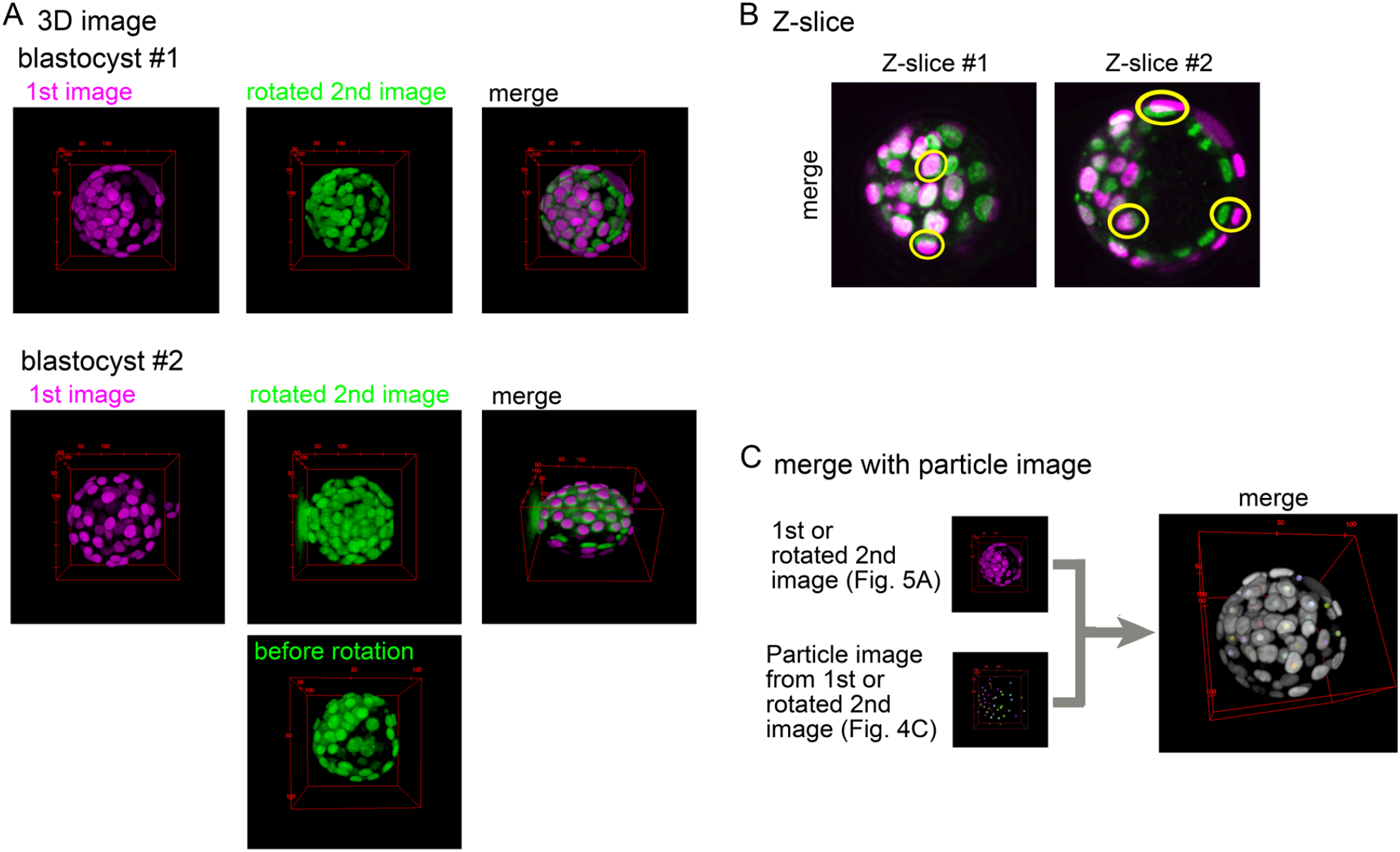
3D reconstruction of rotated image A. 3D images of the 1^st^ and the rotated 2^nd^ image are shown for two blastocysts (#1 and #2). The 2^nd^ images before rotation are shown in Fig. 2 for #1 or in the bottom panel for #2. Note that before the rotation, the intensities of the 2^nd^ images were normalized along the Z-axis (ImageJ>Process>Enhance Contrast…>Normalize), and thus, the intensities were not conserved. B. Two z-slices of the merged image of the blastocyst #1 are exemplified. Yellow, examples of paired nuclei between the 1^st^ and the rotated 2^nd^ images. C. A merged image is exemplified where images of nuclei can be the 1^st^ or the rotated 2^nd^ images, and particles images constructed in Fig. 4 can be the 1^st^ or the rotated 2^nd^ images. In other words, 4 (2×2) combinations of merged images can be generated. The merged image was generated by ImageJ>Image>Color>Merge Channels…

As an additional function in our tool, we can generate merged images between the nuclear images and the particle images (Fig. 5C), which is useful to identify the IDs of the nuclei in the nuclear images.

### Performance and accuracy of our algorithm

We evaluated the performance of our algorithm. In our method, we searched for the optimal values of the three rotation angles as described previously. This kind of problem is called a minimization problem. In general, a minimization problem has a risk that the outcome is trapped at local minima of the cost function to be minimized but not the global minimum which provides the optimal values. In our case, the cost function is the summation of distances between the paired landmarks as defined in Materials & Methods.

To reduce the risk, we performed multiple sets of minimizations in parallel from different initial values of the three rotation angles. For each angle, we set three initial values, resulting in 27 (3×3×3) sets of minimization processes running (Fig. 6A, vertical axis, “27 trials”). Some sets may reach at the global minimum, while other sets may be trapped at local minima. In Fig. 6A, we calculated the probability of reaching the global minimum (i.e. the numbers of the trials reaching the global minimum among the 27 trials). In the case of landmarks = 9, three blastocysts showed high probabilities (#2,3,4), whereas one blastocyst showed low probability (#1). Among the 9 landmarks, we randomly selected 3, 5, or 7 landmarks, and performed the minimizations. For each blastocyst, the number of the landmarks did not significantly affect the probability. These results suggest that the probability is largely dependent on individual blastocysts but not the numbers of landmarks.

**Figure 6:**
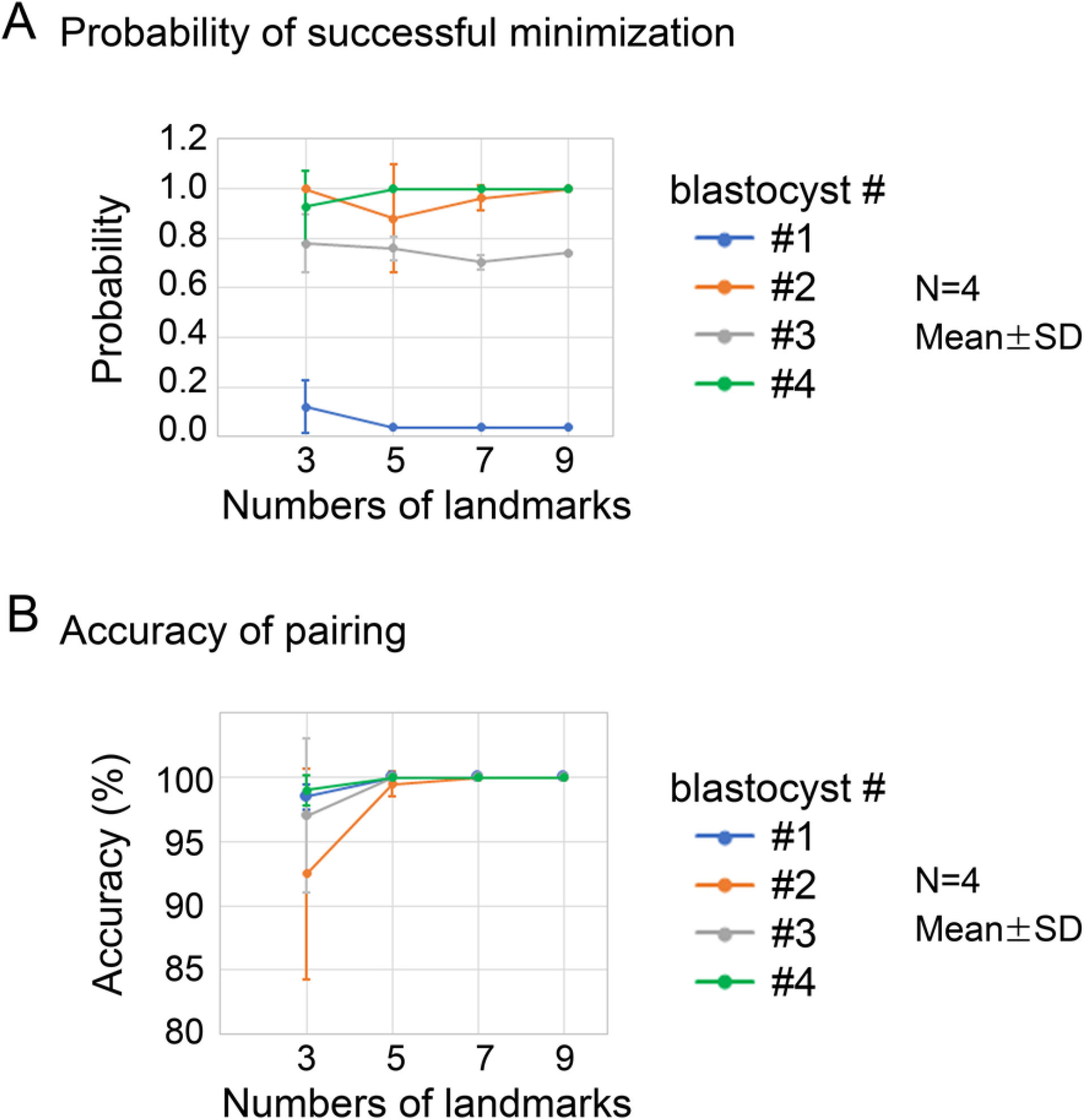
Performance and accuracy of 3D registration A. The performance of the minimization process was evaluated. The probability of successful minimization among 27 trials is shown for each blastocyst (#1-#4); the probability = 1.0 means that all 27 trials successfully reached the global minimum. For the numbers of landmarks = 9, *N* = 1. For the numbers = 3, 5, or 7, landmarks were randomly chosen from the 9 landmarks, and 4 sets of different landmarks were generated; *N* = 4. B. Accuracy of pairing of objects was evaluated for the outcomes of the successful minimization in A. Similar to A, the 4 blastocysts were tested with different number of landmarks for each blastocyst.

Next, we evaluated the accuracy of pairing the nuclei other than the landmarks. We manually defined the correct pairs of the nuclei, and examined whether the outcomes of the minimization are consistent with the correct pairs. Note that we only considered the outcomes of the global minimum. Fig. 6B shows the percentage of the correct pairs of the nuclei. In the case of landmarks = 9, all pairs obtained by the minimization were correct (i.e. accuracy = 100%). On the other hand, under smaller numbers of landmarks (3 and 5), the accuracy became reduced. Together with Fig. 6A, we think that the numbers of landmarks should be ≥ 7, and that 27 sets of initial values of the three rotation angles are sufficient for most of samples.

## Discussion

In the present study, we developed a landmark-based 3D registration tool with subsequent 3D image reconstruction, and demonstrated that this tool worked well for the mouse blastocysts composed of several tens of nuclei. This tool contains several ways of visualizing and quantifying the registration outcomes, which enables us to objectively judge which nucleus in one image corresponds to a nucleus in another image. Importantly, for versatility in the field of experimental biology, this tool can run as ImageJ’s macros.

### Applicability to objects other than nucleus

We chose nuclei as landmarks, but any objects are permitted. A sole requirement of our tool is that users can identify the same position between the 1^st^ and 2^nd^ images. Therefore, even in the case that different markers are used between the two images, we can perform 3D registration if landmarks are correctly defined. Similarly, objects of interest (Fig. 3, step-2) are not limited to nuclei. In addition, even if we do not choose any objects of interest, we can carry out 3D image reconstruction using landmarks. These flexibilities of our tool expand the range of the applicability.

### Comparison with other possible methods

Here we discuss the comparison of our method with other 3D registration methods. Except for the methods described in the Introduction section, other possible method is as follows. The most straightforward strategy of 3D registration is based on pixel-by-pixel correlation of intensities between two images. By translating and rotating one of the two images, we can search for the image transformation which gives the highest correlation. In order that this method works well, there would be some requirements. For instance, decay of intensities along Z-depth should be slight, because the decay significantly affects the value of the correlation. However, in real images of biological specimen, intensities are usually decayed along the Z-depth. Another requirement is related to image qualities, but we cannot expect comparable qualities between images from live specimen and from fixed specimen. A more critical point is its limited applicability to the cases that two specimens are stained by different markers showing different localizations: nucleus vs cytoplasm, something fluorescently labeled vs micro-CT, etc. In these cases, the correlation of intensities between the two images is meaningless. From the viewpoint of computational load, the search for the highest correlation in 3D translation and rotation may result in unrealistic running time. By contrast, our method is based on the manual choice of landmarks. Although this strategy is primitive, we can potentially label correct landmarks under the above situations by considering information of tissue geometries, etc. Therefore, we think that the manual choice of landmarks expands the applicability of our method.

### Expertise required to implement the method

Our method was developed as ImageJ/Fiji’s macros (Note that we recommend Fiji but not ImageJ.). For computers where Fiji is installed, the macros can run immediately after downloading them (i.e. no additional setting). Usual laptops are sufficient to run the macros (e.g. MacBook Air). We provide the protocols with the macros as supplementary materials.

## Supporting information

Protocol of our tools

## Appendix

### Definition of 3D rotation

In the case of 2-dimensional situations, the rotation is defined as the following matrix; 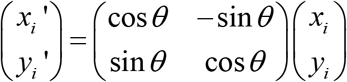, where *θ* is the rotation angle, *x* and *y* are the original coordinates, *x’* and *y’* are the coordinates after rotation, and *i* is the position of *i*th object such as nuclei and pixels. In the case of 3-dimensional situations, the rotation matrix is defined as follows;

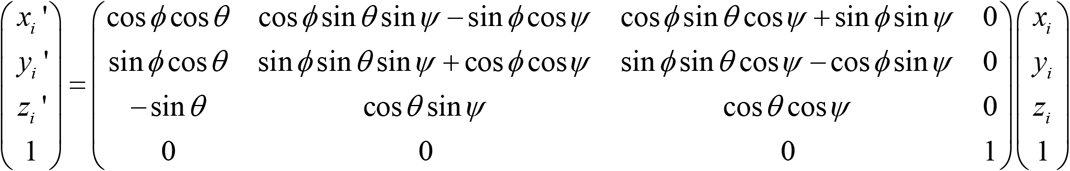

where *ϕ, θ*, and *Ψ* are the angles expressing the three-dimensional rotations.

### Cost function to be minimized

To fit the xyz-coordinates of landmarks from a 2^nd^ image to those from a 1^st^ image by 3D rotation, we considered the summation of the distances between paired landmarks from the two images as follows; 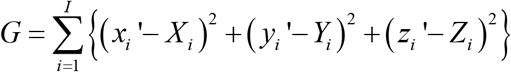, where *G* is the summation, *I* is the total number of the landmarks, and *X, Y*, and *Z* are the xyz-coordinates of the landmarks from the 1^st^ image. We searched for the values of *ϕ, θ*, and *Ψ* which minimized the value of *G*, and thus, *G* is the cost function to be minimized.

### Minimization procedure

The minimization of *G* was achieved using a Monte Carlo algorithm. Initially, *ϕ, θ*, and *Ψ* were set to be certain values: in the case of Fig. 6 where 27 sets of initial values were provided, the values of each angle were 0, (2/3) π, or (4/3) π. These values were iteratively modified so that the value of *G* became smaller, and consequently, *G* is expected to be minimized (i.e. reaches the global minimum) unless the process is trapped at local minima. The minimization procedure was implemented as a macro of ImageJ (Macro_3D_particle_registration_06_v2.ijm).

### 3D depiction of landmarks and objects of interest

According to the xyz-coordinated of landmarks and objects of interest, they were depicted as particles in 3D image (i.e. multiple Z-slices) (Fig. 4). This was implemented as a macro of ImageJ (Macro_particle_drawing_02.ijm). The radius of the particles can be set by users.

### 3D image reconstruction

When we reconstructed 3D rotated image, we applied the values of the three angles to the xyz-coordinates of each pixel/voxel. The intensities of the rotated voxels are averaged by a mean filter of 3 × 3 × 3 kernel. This was implemented as a macro of ImageJ (Macro_3D_image_rotation_02.ijm).

