## Supplementary material for "An ImageJ-based tool for three-dimensional registration between different types of microscopic images": Protocol of our tools

### Protocol of 3D registration

Title:

Koyama et al.

Our method is composed of the following three macros. For each macro opened, users can find the essential information; descriptions, requirements, setting of parameter values to be done by users, tips, etc. Only the most fundamental processes and parameter settings are explained in this presentation.

Macro_3D_particle_registration_06_v2.ijm

Macro_particle_drawing_02.ijm

Macro_3D_image_rotation_02.ijm

### Macro name: Macro_3D_particle_registration_06_v2.ijm

#### Format of input file

File name should be “input_xyz_registration.csv”

(Csv files are generated by ImageJ>”Multi-point tool” followed by “Analyze>Measure”, and then edited on the Excel software.)

x | y | z | From 1st or 2nd image | ID

| | A | B | C | D | E |
|---|---|---|---|---|---|
| 1 | 42.813 | 12.706 | 39.1 | 1 | 1 |
| 2 | 34.665 | 18.921 | 26.45 | 1 | 2 |
| 3 | 72.92 | 74.025 | 10.925 | 1 | 3 |
| 4 | 50.547 | 86.869 | 12.65 | 1 | 4 |
| 5 | 33.836 | 89.355 | 29.325 | 1 | 5 |
| 6 | 15.744 | 70.573 | 31.05 | 1 | 6 |
| 7 | 21.821 | 31.074 | 35.65 | 1 | 7 |
| 8 | 21.821 | 58.833 | 14.375 | 1 | 8 |
| 9 | 18.23 | 45.023 | 20.7 | 1 | 9 |
| 10 | 28.588 | 37.703 | 11.5 | 1 | 10 |
| 11 | 65.325 | 26.24 | 13.8 | 1 | 11 |
| 12 | 81.207 | 47.923 | 13.225 | 1 | 12 |
| 13 | 85.902 | 67.81 | 14.95 | 1 | 13 |
| | ,,, | ,,, | ,,, | ,,, | ,,, |
| | ,,, | ,,, | ,,, | ,,, | ,,, |

Paired landmarks should have the same ID.

Objects of interest should have the different IDs.

| | A | B | C | D | E |
|---|---|---|---|---|---|
| 43 | 25.135 | 62.286 | 31.05 | 1 | 43 |
| 44 | 33.146 | 64.496 | 27.025 | 1 | 44 |
| 45 | 39.913 | 73.887 | 27.025 | 1 | 45 |
| 46 | 31.35 | 68.501 | 9.2 | 1 | 46 |
| 47 | 52.619 | 12.015 | 45.625 | 2 | 1 |
| 48 | 43.227 | 13.258 | 33.125 | 2 | 2 |
| 49 | 59.8 | 66.844 | 3.75 | 2 | 3 |
| 50 | 35.632 | 73.058 | 7.5 | 2 | 4 |
| 51 | 20.578 | 72.506 | 21.25 | 2 | 5 |
| 52 | 11.049 | 52.757 | 29.375 | 2 | 6 |
| 53 | 28.45 | 21.821 | 43.75 | 2 | 7 |
| 54 | 20.025 | 41.57 | 16.875 | 2 | 8 |
| 55 | 20.302 | 29.831 | 25.625 | 2 | 9 |
| 56 | 31.903 | 26.102 | 16.875 | 2 | 47 |
| 57 | 67.396 | 25.964 | 17.5 | 2 | 48 |
| 58 | 74.025 | 46.956 | 9.375 | 2 | 49 |
| | ,,, | ,,, | ,,, | ,,, | ,,, |
| | ,,, | ,,, | ,,, | ,,, | ,,, |

### Macro name: Macro_3D_particle_registration_06_v2.ijm

#### Run of the macro

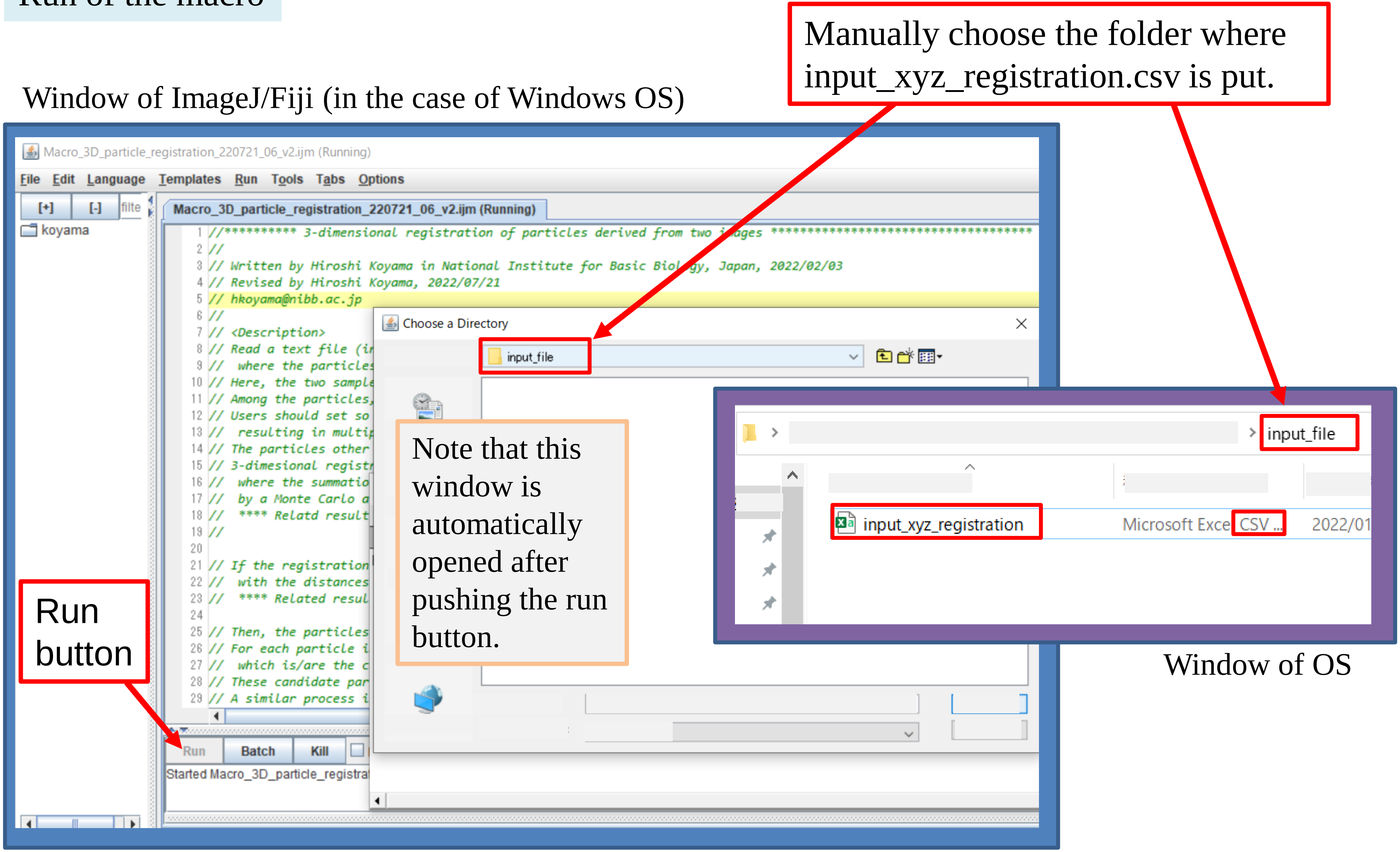

### Macro name: Macro_3D_particle_registration_06_v2.ijm

#### Output files of the macro

- input_xyz_registration
- Log_registration → Log file
- output_bad_results_rotation → Unsuccessful minimization
- output_condition_for_registration → Parameter values/conditions
- output_registered_particles_1neighbors → List of candidates: 1 nearest neighbor
- output_registered_particles_3neighbors → List of candidates: 1-3 nearest neighbors (Fig. 4C)
- output_registered_particles_all → IDs and xyz-coordinates of paired landmarks/objects between the 1st and the rotated 2nd images (Fig. 4B,D)
- output_registered_particles_landmarks → IDs and xyz-coordinates of paired landmarks between the 1st and the rotated 2nd images (Fig. 4A)
- output_rotation_results → List of successful minimization with the values of the three angles, and the best values (Fig. 5)

List of successful minimization

```
1: 9636 xyz-error(2357.9223 -> 112.7069) mean-distance(2.6259, 3.4258) angle(5.9524, 6.229, 0.2563)
,,,
best rotational angles: 5.9524, 6.229, 0.2563
```

The best values

Macro name: Macro_3D_particle_registration_06_v2.ijm

A format of an output file (output_registered_particles_3neighbor.txt, related to Fig. 4C)

At the end of the text file in Fig. 4C, multiply assigned objects are listed as follows.

,,, ,,, ,,,
,,, ,,, ,,,
,,, 61 ,,,
,,, ,,, ,,,
,,, ,,, ,,,

| | A | B | C | D | E | F | G |
|---|---|---|---|---|---|---|---|
| 38 | 45 | 82 | 72 | 61 | 5.46 | 12.108 | 12.215 |
| 39 | 46 | 61 | 56 | 62 | 17.083 | 17.87 | 19.572 |
| 40 | multiply counted particles = 2 | | | | | | |
| 41 | 55 61 | | | | | | |

See Fig. 4C

This means that ID = #55 and #61 in the 2nd image are multiply assigned as the nearest neighbor.

### Macro name: Macro_3D_particle_registration_06_v2.ijm

#### Optional for correction of distortion of xyz-coordinates: step-3 in Fig. 3A

Window of the macro on ImageJ/Fiji

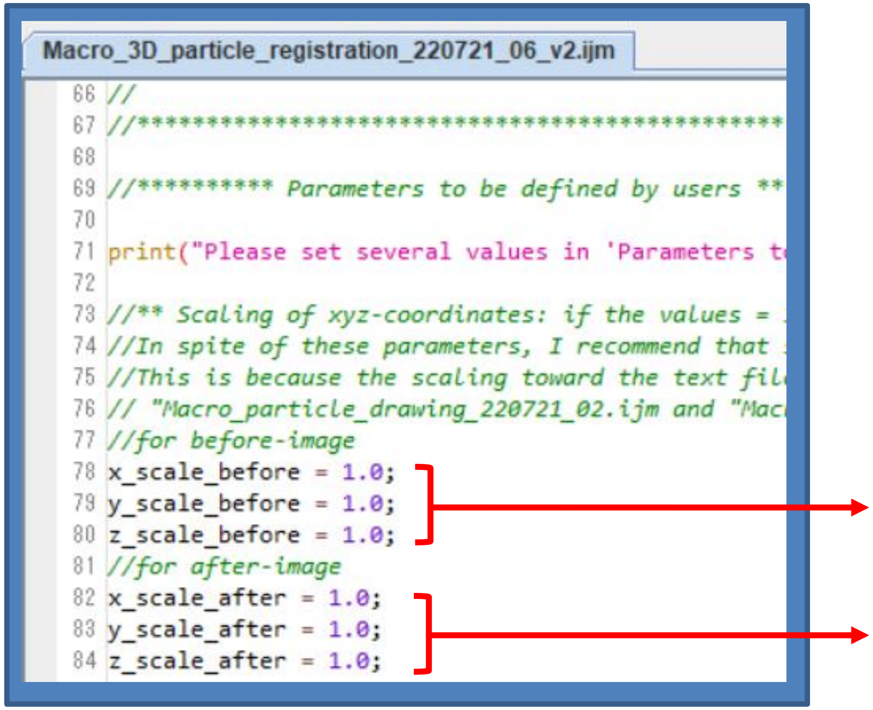


Enter manually the magnification values for xyz-coordinates in the 1st image

Enter manually the magnification values for xyz-coordinates in the 2nd image

Before running the macro, users rewrite these values.

Important!
Note that these values will be used in the following two macros, "Macro_particle_drawing_02.ijm" and "Macro_3D_image_rotation_02.ijm", where users should manually rewrite the corresponding lines in the two macros. Otherwise, images with different xyz-scales are generated.

### Macro name: Macro_particle_drawing_02.ijm

#### Parameter setting

Window of the macro on ImageJ/Fiji

Macro_particle_drawing_220721_02.ijm

```


//****** Basic parameters to be defined by users ******

print("Please set several values in 'Parameters to be defined by us

//Choose input text file
input_name = "output_registered_particles_all.txt";
//input_name = "output_registered_particles_landmarks.txt";

image_size_x = 262;     //The pixel number of output images along x
image_size_y = 262;     //The pixel number of output images along y

//Unit of input text file
//If the unit is pixel, um_pixel should be set as 1.0.
//If the unit is um or something, um_pixel should be set so that th
///The relationship between um and pixel in the orignal before-imag
um_per_pixel = 0.4143204;

//These values should be the same as those used in Macro_3D_particl
x_scale_before = 1.0;
y_scale_before = 1.0;

radius_of_particle=5.0;              //Particle radius to be drawn
```

Users should manually write the following parameters.

Choose one of two possible input files obtained from the previous macro.

Image sizes

Unit of length: µm/pixel

See the previous slide.

Size of particles to be drawn

### Macro name: Macro_3D_image_rotation_02.ijm

#### Parameter setting

Window of the macro on ImageJ/Fiji

Macro_3D_image_rotation_220722_02.ijm

```
//** Rotation angles for after-image, which should be
//output_rotation_results.txt after running the Macro_
//The values are written in the last line of the text
angle1=5.9529;
angle2=6.2313;
angle3=0.2541;

//** Nomalization of intensity gradient along z-slices
//If your images exhibit severe decay of intensity in
z_normalize = 1;    //1 for Yes, 0 for No.

//** Averaging of 3D image drawing
//If range_ave = 0, no averaging. If range_ave = 1, 3x
// This averaging improves image quality, but time con
range_ave = 1;

//*****************************************************


//***** input text file name and input after-image nam
dir_input = getDirectory("Choose a Directory ");
input_file = dir_input+"input_xyz_registration.csv";
input_image = dir_input+"input_after_image.tif";
```

Users should manually write the following parameters.

The values of the three angles obtained from the 1st macro.

Before rotation, the intensities of the 2nd image is normalized or not.

Input file also used in the 1st macro

The name of the original 2nd image to be rotated.

Note that the parameters of xyz-scaling are also to be set as previously explained.
